# Protracted fate acquisition and epigenetic de-aging during induced neural stem cell conversion of human blood cells

**DOI:** 10.1101/2025.10.25.683217

**Authors:** Lea Jessica Berg, Julia Franzen, Andreas Buness, Rachel Konang, Chao Sheng, Michael Peitz, Andreas Till, Wolfgang Wagner, Oliver Brüstle

**Author notes:** Genomics Facility, University Hospital Aachen, Pauwelsstraße 30, 52074 Aachen, Germany (Julia Franzen). CCRM Nordic AB, GoCo House, Förändringens Gata 10, 431 53 Mölndal, Sweden. Federal Institute for Drugs and Medical Devices (BfArM), Kurt-Georg-Kiesinger-Allee 3, 53175 Bonn, Germany (Andreas Till).

## Abstract

Transcription factor-based direct conversion of somatic cells represents an interesting avenue for generating induced neural stem cells (iNSCs) from peripheral blood without transit through a pluripotent stage. While this paradigm has been shown to be associated with epigenetic de-aging, the dynamics of this process have remained unclear. Here, we used overexpression of the two reprogramming factors SOX2 and cMYC to generate iNSC from erythroid progenitors of donors ranging from neonatal to 101 years of age. Using an epigenetic clock algorithm, we corroborated our previous finding that iNSCs generated from aged donors show pronounced epigenetic de-aging, preserving around 13 % and 5 % of the original donor age at low and high passages, respectively. Studying the dynamic of epigenetic de-aging during iNSC conversion across time, we found that this process is largely protracted, continuing for several weeks and even beyond forced neuronal differentiation of iNSCs. Transcriptomic differences between young and old donor-derived iNSCs dissipate with extended time in conversion, too. Concordant with this observation, established iNSC lines lack age-associated cellular hallmarks, similar to induced pluripotent stem cells and their derivatives. Interestingly, time course analysis of DNA methylation and RNA sequencing data revealed that acquisition of a *bona fide* NSC signature extends greatly beyond the time point when proliferative PAX6-positive iNSCs emerge. The unexpected slow dynamics of these processes makes iNSC conversion an attractive model for dissecting the mechanisms underlying somatic transdifferentiation and de-aging.

**Statement of Ethical Approvals:** The collection of human somatic material (*i.e.*, blood) from healthy donors for iPSC reprogramming and iNSC conversion was conducted in accordance with German law and approved by the Ethics Committee of the University of Bonn Medical Center (approval number: 275/08). All subjects gave written informed consent.

## Introduction

The rise of cell programming, encompassing all techniques that aim at manipulating cellular identity such as induced pluripotent stem cell (iPSC) reprogramming as well as direct cell fate conversion, has largely facilitated the derivation of human neural cells for applications such as disease modeling and regenerative cell replacement. Significant remodeling events of transcriptomic and epigenetic signatures play a pivotal role in the process of cell programming. Yet, the dynamics and underlying mechanisms of these processes are only partially understood. During iPSC reprogramming, genome-wide remodeling is induced, first leading to the silencing of the cell of origin’s transcriptional program, followed by the activation of transient enhancers and eventually upregulation of pluripotency-associated genes such as *Sall4, Nanog* and *Dppa2* that initiate, mature and/or stabilize the acquired pluripotent state (Buganim, Faddah, & Jaenisch, 2013; Deng, Jacobson, Collier, & Plath, 2021). Direct conversion into post-mitotic induced neurons (iN), which was first successfully achieved by the overexpression of the transcription factor cocktail Ascl1, Brn2 plus Myt1l in mouse fibroblasts (Vierbuchen et al., 2010), offers a more direct route for generating neural cells without a pluripotent intermediate.

Interestingly, iPSC reprogramming and direct conversion into iNs have vastly diverging effects on cellular aging signatures: Cellular age, if assessed using DNA methylation (DNAm)-based algorithms (Horvath, 2013; Horvath et al., 2018), has been shown to be reverted to an embryonic stage upon iPSC reprogramming (Horvath, 2013; Lo Sardo et al., 2017; Weidner et al., 2014), while iNs retain the DNAm age of their source cells (Huh et al., 2016). Differential preservation of age-associated cellular traits between iPSCs and iNs is also reflected by the length of telomeres, mitochondrial health, nuclear organization and the extent of transcriptional senescence signatures (Kim et al., 2018; Lapasset et al., 2011; Mertens et al., 2015; Miller et al., 2013; Prigione et al., 2011; Tang, Liu, Zang, & Zhang, 2017; Yang et al., 2015).

In a previous study (Sheng et al., 2018), we have shown that human PBMC-derived erythroid progenitor cells (EPCs) can be converted into clonally growing iNSCs by SeV-mediated overexpression of the two transcription factors SOX2 and cMYC. iNSCs exhibit tripotent differentiation capacity, with neuronal differentiation being amenable to regional patterning, and can be employed for disease modeling and neurotransplantation. While we could further demonstrate that iNSCs are overall similar to iPSC-derived NSCs, not only with regard to transcriptomic, but also epigenetic and cellular age-associated signatures (Sheng et al., 2018), the question how signatures of cellular identity and age are remodeled in the process of conversion was not yet addressed. Here, we set out to explore the dynamics of cell fate conversion on the one hand and loss of aging signatures on the other. To this end, we generated and characterized isogenic pairs of iNSCs and iPSC-derived NSCs from blood of newborns to high-age donors, representing the entire human life span. We then dissected the blood-to-iNSC conversion process by performing DNAm and RNA sequencing analyses across the time course of transdifferentiation. To our surprise, our findings demonstrate that both processes, epigenetic de-aging and cell fate consolidation, are largely protracted, with epigenetic de-aging even extending into the postmitotic neuronal stage.

## Methods

### Cell culture techniques EPC induction

Peripheral blood mononuclear cells (PBMCs) were isolated from fresh blood draws using vacutainer CPT tubes (BD Bioscience, San Jose, USA), which were centrifuged for 30 minutes at 1,800 rcf. Isolated mononuclear cells were washed twice with 1x DPBS (Thermo Fisher Scientific, Waltham, USA) before 3-5×10^6^ PBMCs were either frozen in 90 % FCS (Thermo Fisher Scientific) plus 10 % DMSO (Sigma-Aldrich, St. Louis, USA) or directly resuspended in EPC medium consisting of StemSpan SFEM (Stemcell Technologies, Vancouver, Canada) supplemented with 1 % CD Lipid (Thermo Fisher Scientific), 2 U/ml recombinant human (rh) erythropoietin (Bio-Techne, Minneapolis, USA), 100 ng/ml rhSCF (Bio-Techne), 40 ng/ml rhIGF1 (Bio-Techne), 1 µM dexamethasone (Sigma-Aldrich) and 10 ng/ml rhIL3 (Bio-Techne) for subsequent EPC induction, which was performed according to a published protocol with minor modifications (van den Akker, Satchwell, Pellegrin, Daniels, & Toye, 2010). In short, cells were cultured in uncoated 6-well cell culture plates at 6 % CO_2_ and 37 °C. From days 1 to 6 of EPC genesis, 1 ml EPC medium was added freshly per day and well. At day 7, EPCs were enriched by applying a Percoll (Sigma-Aldrich) density gradient with a density of 1.075 ρ. After centrifugation, the interphase ring containing EPCs was collected. Cells were washed twice before EPCs were resuspended in EPC medium lacking rhIL3 and plated in uncoated 6-well cell culture-plates at a density of 1-1.5×10^6^ cells/6-well for further maturation. Enriched EPCs were expanded until day 9 of EPC genesis, performing medium addition on day 8. On day 9, EPC-containing cell suspensions were collected and 1-2×10^6^ cells were frozen in 90 % EPC medium lacking rhIL3 but supplemented with 10 % DMSO.

### iPSC reprogramming and differentiation into NSCs

EPCs were thawed in DMEM/F12 (Thermo Fisher Scientific). After 1-hour incubation at 6 % CO_2_ and 37 °C, 3×10^5^ EPCs were aliquoted in 15 ml-tubes and centrifuged for 5 minutes at 1,500 rpm. After removing the supernatants, cell pellets were resuspended in 250 µl EPC medium without rhIL3 containing the CytoTune 2.0 (Thermo Fisher Scientific) Sendai viruses (SeVs) – *i.e.*, a polycistronic vector for KLF4-OCT3/4-SOX2 overexpression, as well as single vectors encoding for cMYC and KLF4 – diluted to a multiplicity of infection (MOI) of 5:5:1. Solutions containing cells and viruses were then transferred to uncoated 4-wells and incubated for approximately 24 hours at 6 % CO_2_ and 37 °C. The next day, infected cells were collected and centrifuged for 5 minutes at 1,500 rpm. The virus-containing supernatants were discarded, cell pellets resuspended in 500 µl EPC medium without rhIL3 and plated in uncoated 4-wells. On day 3 of iPSC reprogramming, cells were collected and centrifuged 5 minutes at 1,300 rpm. Supernatants were discarded and cells were replated in EPC medium without rhIL3 on 10 cm cell culture-dishes coated with Matrigel (Corning Inc., Corning, USA; 1:60 dilution in DMEM/F12). Medium was gradually changed to StemMACS^TM^ iPS-Brew XF (incl. supplement; Miltenyi Biotec, Bergisch Gladbach, Germany) from days 5 to 11 of iPSC reprogramming by performing partial medium changes with increasing concentrations of StemMACS^TM^ iPS-Brew XF. Around day 20 of iPSC reprogramming, up to 24 colonies were manually picked under microscopic control and plated in StemMACS^TM^ iPS-Brew XF supplemented with 10 µM Rock-inhibitor Y-27632 (Cell Guidance Systems, Cambridge, UK) onto Matrigel-coated 24-well plates (1:60 dilution in DMEM/F12). In order to eliminate SeV transgenes, this procedure was repeated every week until five single colony picking cycles were completed. After the fifth picking cycle, cells were allowed to grow confluent. Once confluency was reached, cells were dissociated by 3-minute-long incubation with 0.5 mM EDTA (Carl Roth, Karlsruhe, Germany). After removing the EDTA, cells were detached by pipetting and replated in cell culture formats appropriate for the number of cells detached. Transgene-free iPSC lines were routinely expanded in StemMACS^TM^ iPS-Brew XF on vitronectin-coated (Thermo Fisher Scientific; 1:100-1:200 dilution in 1x DPBS) cell culture formats before being frozen in 90 % StemMACS^TM^ iPS-Brew XF plus 10 % DMSO for cryo-preservation.

In order to generate a stable NSC population from iPSCs, a protocol adapted after Reinhardt *et al*. (2013) was applied. First, iPSCs were singularized by 5-15 minutes-long incubation with StemPro^TM^ Accutase^TM^ (Thermo Fisher Scientific) at 37 °C. Cell suspensions were diluted in DMEM/F12 and centrifuged for 5 minutes at 1,000 rpm. Afterwards, 2×10^6^ iPSCs were resuspended in 1xN2B27 (1:1 DMEM/F12 : Neurobasal with 0.5x N2 supplement, 0.5x B27 supplement without vitamin A, 2 mM L-glutamine, 0.0025% BSA and 1x Pen/Strep; all materials from Thermo Fisher Scientific) containing 3 µM CHIR99021 (Miltenyi Biotec), 0.5 µM purmorphamine (Miltenyi Biotec), 10 µM SB431542 (Axon Biotech, Hengersberg, Germany) and 1 µM dorsomorphine (Tocris Bioscience, Bristol, UK) and supplemented with 10 µM Y-27632, and transferred to AggreWell^TM^ 800 microwell culture plates (Stemcell Technologies), which were prepared according to the manufacturer’s instructions. AggreWell^TM^ 800 plates were centrifuged for 3 minutes at 300 rpm. The next day, a half medium change with 1xN2B27 containing CHIR99021, purmorphamine, SB431542 and dorsomorphine was performed. Approximately 48 hours after aggregation of iPSCs to embryoid bodies, these were carefully dislodged from the AggreWells and transferred to uncoated 6 cm Petri-dishes. One day later, medium was changed to 1xN2B27 medium supplemented with 3 µM CHIR99021, 0.5 µM purmorphamine and 64 µg/ml LAAP (Sigma-Aldrich)). After another medium change on day 4 of NSC generation, embryoid bodies were triturated in small pieces on day 5 using a 1 ml-pipette. Triturated embryoid bodies from one AggreWell were distributed to a full Matrigel-coated (1:60 dilution in DMEM/F12) 6-well cell culture-plate and seeded in 1xN2B27+Pen/Strep+CPL supplemented with 10 µM Y-27632. From this day onwards, NSCs were routinely cultivated on Matrigel-coated cell culture formats (1:60 dilution in DMEM/F12) in medium without Pen/Strep. Medium was changed every 1 to 2 days, and cells were passaged when reaching confluency using Accutase. Splitting ratios typically ranged from 1:4 - 1:12. Up until passage 5, 10 µM Y-27632 was added for replating.

### iNSC conversion and propagation

EPCs were converted into iNSCs according to the procedure published in Sheng *et al*. (2018) with minor modifications. On day 0 of iNSC conversion, 1.5×10^5^ EPCs were thawed as described and spin-infected in EPC medium without rhIL3 containing SeV-SOX2 and SeV-cMYC (each at MOI 5, both single vector-SeVs included in the CytoTune 1.0 kit (Thermo Fisher Scientific, now purchased from ID Pharma, Tokyo, Japan) for 30 minutes at 1,500 rcf and 32 °C. Cell pellets were resuspended in their same virus-containing supernatant after centrifugation and plated into uncoated 4-wells. Cells were kept in a humidified incubator for approximately 24 hours at 37 °C and 21 % O_2_. The next day, suspension cells were collected, pelleted by 5-minute-long centrifugation at 1,500 rpm, resuspended in 75 % iNSC conversion medium, consisting of 2x N2B27 (1:1 Advanced DMEM/F12 (Thermo Fisher Scientific) : Neurobasal with 1x N2 supplement, 1x B27 supplement without vitamin A, 2 mM L-glutamine and 0.0025% BSA) supplemented with 3 µM CHIR99021, 1 µM purmorphamine, 0.5 µM A83-01 (Tocris Bioscience), 10 ng/ml human LIF (Novoprotein, Fremont, USA), 64 µg/ml LAAP (Sigma-Aldrich), 1 µg/ml laminin (Sigma-Aldrich) and 5 µM tranylcypromine (Enzo Biochem Inc., Farmingdale, USA), plus 25 % EPC medium without rhIL3 and plated onto Matrigel-coated cell culture formats (1:60 dilution in DMEM/F12) at a density of about 1.3×10^4^ cells/cm^2^. From this day onwards, converting cells were cultured at 37 °C and 5 % O_2_. On day 3 of conversion, medium consisting of 75 % iNSC conversion medium plus 25 % EPC medium without rhIL3 was added. From day 5 onwards, emerging adherent iNSCs were cultured in 100 % iNSC conversion medium, performing full medium changes every other day. Starting from day 10 to 11, iNSCs were cultivated in iNSC expansion medium (2x N2B27 supplemented with 3 µM CHIR99021, 0.5 µM purmorphamine, 0.5 µM A83-01, 10 ng/ml human LIF and 64 µg/ml LAAP). Colonies consisting of neuroepithelial-like shaped cells were picked during the first three weeks of iNSC conversion under microscopic control as described. In order to generate polyclonal cell lines, remaining colonies were dissociated on day 21 of conversion with Accutase as described, pooled and plated in Matrigel-coated cell culture formats (1:60 dilution in DMEM/F12) at a density of approximately 1×10^5^ cells/cm^2^. Medium of iNSC lines was changed every 1 to 2 days. Lines were passaged at least once a week, ideally when cells had reached a confluency > 100 %, using Accutase. Typical splitting ratios ranged from 1:3 - 1:9. From day 35 of conversion onwards, iNSCs were cultured under normoxic conditions at 37 °C and 21 % O_2_. SeV removal was achieved by prolonged cultivation (typically between 4 and 7 weeks) of established iNSC lines at 39 °C. During this period, SeV-mediated transgene expression was monitored weekly by RT-PCR. Once expression of both SeVs was below the detection level of the RT-PCR, cells were transferred back to 37 °C. One week thereafter, another RT-PCR was performed in order to confirm the persistent absence of SeVs.

For manipulating proliferation speed during EPC-to-iNSC conversion, cells were treated with either 2 % (v/v) glycerol (Sigma-Aldrich) or 2 mM thymidine (Sigma-Aldrich) starting from day 14 of conversion. During this time, iNSCs were replated once a week and cell numbers were estimated at each replating step using a Neubauer counting chamber (Paul Marienfeld, Lauda-Königshofen, Germany) in order to calculate population doublings. Glycerol was added to the medium with every medium change but was omitted on days when splitting was performed. Thymidine treatment always started the day after splitting and was discontinued after 48 hours and 24 hours for weeks 2 to 5 and 6 to 7 of conversion, respectively.

### iNSC differentiation by NGN2 overexpression

A transcription factor-mediated forward programming approach was implemented for rapid differentiation of iNSCs into post-mitotic neurons early in the conversion process. To this end, at day 7 of conversion, iNSCs were infected with two lentiviruses (1:50-1:80 dilution), encoding for a reverse tetracycline-controlled transactivator (rtTA) and a tetracycline-responsive promoter element (TRE)-regulated NGN2 expression cassette. Afterwards, iNSCs were routinely cultured up until day 21 of conversion, when iNSCs were replated at a density of 8.3×10^4^ cells/cm^2^. The day after, forward programming was induced by changing medium to NGMC containing 10 ng/ml BDNF, 10 ng/ml GDNF, 64 µg/ml LAAP, 5 µM DAPT and 1 µg/ml DOX. Differentiation of iNSCs was proceeded under normoxic conditions at 37 °C and 21 % O_2_. From day 5 to 7 of differentiation, 5 µM Cytarabin/Ara-C (Merck Millipore) was added to the differentiation medium in order to kill mitotically active cells. Medium was changed three times a week during the 20-day-long differentiation period.

### Molecular biology techniques DNAm analysis

DNA extraction was performed using Qiagen’s (Hilden, Germany) Blood & Tissue kit according to the manufacturer’s instructions. The concentration and purity of extracted DNA were determined using a Nanodrop 2000c. For DNAm profiling, DNA samples were diluted to a concentration of 55 ng/µl with AE buffer (Qiagen) and provided to the Next Generation Sequencing Core Facility of the University of Bonn. DNA samples were then bisulfite-converted before being profiled on Infinium MethylationEPIC 850k chips (Illumina). Raw data were processed using the minfi package (Aryee et al., 2014) in the R environment (R Core Team, 2019. R: A Language and Environment for Statistical Computing. R Foundation for Statistical Computing, Vienna, Austria. https://www.R-project.org/). Afterwards, retrieved beta values were used to calculate the samples’ epigenetic age (also called DNAm age) using a predictor algorithm based on (Horvath, 2013; Horvath et al., 2018). In addition, DNAm data were assessed for differentially methylated probes and regions (DMPs and DMRs, respectively) in R using ChAMP (Morris et al., 2014; Tian et al., 2017). ChAMP’s gene set enrichment analysis was followed up with rrvgo (Sayols, 2020), and Venn diagrams were plotted using an online tool provided by the Ghent university (http://bioinformatics.psb.ugent.be/webtools/Venn/). In addition to retrieving gene ontology (GO) terms within ChAMP, significant most variable positions (MVPs) identified by this package were mapped to genes using the missMethyl package (Phipson, Maksimovic, & Oshlack, 2016), prior to pathway analysis using Reactome PA (Yu & He, 2016) in R.

### RNA sequencing analysis

RNA extraction was performed with Qiagen’s RNeasy kit according to the manufacturer’s instructions. The concentration and purity of RNA samples were analyzed with a Nanodrop 2000c. RNA samples were stored at −80 °C. For bulk RNA sequencing, RNAs were diluted to a concentration of around 50 ng/µl with RNA-free H_2_O (Qiagen) and provided to the Next Generation Sequencing Core Facility of the University of Bonn, where samples were subjected to QuantSeq 3’mRNA-Seq library preparation (Lexogen GmbH, Vienna, Austria) before being sequenced on a HiSeq 2500 V4 (Illumina) in high output mode.

Raw data were processed by the Bioinformatics Core Facility of the University of Bonn. In short, after trimming of the Illumina Universal Adapter with cutadapt (Martin, 2011), reads have been aligned to the human genome (GRCh38) with STAR (Dobin et al., 2013). featureCounts (Liao, Smyth, & Shi, 2014) was used to assign reads to genes as defined by Ensembl. A read is counted, firstly, if it is uniquely mapped, secondly, if it matches the strand of the gene, and thirdly, if it overlaps with only one gene, *i.e.*, the read can be non-ambiguously assigned to a single gene. Only genes with at least 100 read counts across samples and at least 2 samples each with a minimum count of 40 were considered in the downstream statistical analysis which was performed in the R environment with the Bioconductor package limma (Huber et al., 2015; Ritchie et al., 2015). As samples from the same donor were measured repeatedly over time and cellular state, donor was considered as a random effect in the statistical model and passed to the function duplicateCorrelation as blocking factor. In addition, sequencing batch and sex were modelled as fixed effects to correct for unwanted variation. Statistical contrasts were calculated for the main fixed effect defined by combining age class, cell type and time in a single variable. The Benjamini-Hochberg method was used to calculate p-values adjusted to multiple testing (false discovery rate, FDR) for each contrast. Data visualisation, such as volcano plots and heatmaps were generated using R-packages ggplot2 (Wickham, 2016) and ComplexHeatmap (Gu, Eils, & Schlesner, 2016), respectively. Pathway enrichment analysis (Fisher test) for differently expressed genes (FDR < 0.05) was performed using the Bioconductor package clusterProfiler (Wu et al., 2021). Only pathways with a set size of at least 10 or at most 500 genes were considered. For some analyses, enriched GO terms were depicted as treemap plots with rrvgo as described.

### Immunocytochemistry and fluorescence imaging

For staining cells within the first two weeks of conversion, both adherent cells and cells in suspension were collected. To this end, the supernatant was harvested and adherent cells were dissociated with Accutase as described. The adherent and non-adherent fractions were pooled and subsequently spun down on slides at 500 g for 7 minutes using appropriate spinning cuvettes. Afterwards, primary mouse monoclonal IgG antibody to CD71 (1:100; BD Pharmingen, Franklin Lakes, USA) in HBSS was incubated for 30 minutes at room temperature. Antibody solution was washed off using HBSS before cells were fixed with 4 % PFA (Thermo Fisher Scientific) for 5 minutes at room temperature. Following another washing step with 1x DPBS containing 1 % BSA, slides were incubated with 1x DPBS containing 1 % BSA and 0.5 % Triton-X100 (Sigma-Aldrich) for 30 minutes at room temperature. Afterwards, cells were stained with rabbit polyclonal IgG antibody to DACH1 (1:100; Proteintech, Planegg-Martinsried, Germany) in 1x DPBS containing 1 % BSA and 0.3 % Triton-X100 for 2 hours at room temperature. After washing in HBSS, secondary antibodies goat anti-mouse IgG (H+L) Alexa Fluor 555 and goat anti-rabbit IgG (H+L) Alexa Fluor 488 (both from Thermo Fisher Scientific) in 1x DPBS containing 1 % BSA and 0.3 % Triton-X100 were incubated for 30 minutes at room temperature. Cells were washed again in 1x DPBS containing 1 % BSA before nuclei were stained with 2 µg/ml 4’,6-Diamidino-2-phenyl-indol-dihydrochloride (DAPI; Sigma-Aldrich) for 5 minutes at room temperature. Finally, slides were washed in 1x DPBS and mounted with cover slips using 133.33 mg/ml Mowiol-488 plus 25 mg/ml DABCO (both from Carl Roth).

All adherently growing cell preparations were fixed with 4 % PFA for 10 minutes at room temperature before being washed with 1x DPBS once. Cells were blocked with 10 % FCS in 1x DPBS for 2 hours at room temperature. Blocking solutions were supplemented with 0.5 % Triton-X100 for cell permeabilization. After blocking, cells were incubated overnight at 4 °C with primary antibodies diluted in 1x DPBS containing 5 % FCS + 0.3 % Triton-X100: chicken polyclonal IgY to MAP2 (1:1,000; Bio-Techne), mouse monoclonal IgG to γH2AX (1:250; Merck Millipore), mouse monoclonal IgG to LMNA/C (1:200; Abcam, Cambridge, UK), mouse monoclonal IgM to O4 (1:500; Bio-Techne), mouse monoclonal IgG to S100β (1:1,000; Sigma-Aldrich), mouse monoclonal IgG to SOX2 (1:100; Bio-Techne), mouse monoclonal IgG to TUBB3 (1:1,000; BioLegend, San Diego, USA), rabbit polyclonal to GFAP (1:1,000; Merck Millipore), rabbit polyclonal IgG to LAP2a (1:250; Abcam), rabbit polyclonal IgG to NES (1:200; Bio-Techne) and rabbit monoclonal IgG to DCX (1:200; Abcam). The next day, cells were washed three times with 1x DPBS + 0.3 % Triton-X100 before secondary antibodies, diluted in 1x DPBS containing + 0.3 % Triton-X100, were incubated for 2 hours at room temperature: goat anti-chicken IgY (H+L) Alexa Fluor 647 conjugate, goat anti-mouse IgG (H+L) Alexa Fluor 488 as well as 555 conjugates, goat anti-mouse IgM (H+L) Alexa Fluor 488 conjugate and goat anti-rabbit IgG (H+L) Alexa Fluor 488 as well as 555 conjugates (all from Thermo Fisher Scientific). After three additional washing steps with 1x DPBS + 0.3 % Triton-X100, 2 µg/ml DAPI was incubated for 5 minutes at room temperature to counterstain nuclei. Cells were finally washed once with 1x DPBS before mounting with a medium containing 133.33 mg/ml Mowiol-488 and 25 mg/ml DABCO. Mounted specimens were dried at least overnight at 4 °C before imaging.

Image acquisition of immunofluorescent stainings was either performed manually using Zeiss’ (Oberkochen, Germany) AxioImager or automatedly utilizing the IN Cell Analyzer 2200 (GE Healthcare Bio-Sciences Corp, Picataway, USA). For qualitative image presentation, color brightness and contrast were either adjusted right after image acquisition with the software associated with the utilized microscope or retrospectively using ImageJ (© National Institutes of Health, Bethesda, USA; version 1.52a Java 1.8.0_181, 64-bit). Quantification of immunofluorescent stainings was performed on unmodified raw data using CellProfiler (Kamentsky et al., 2011); © Broad Institute, Cambridge, USA; version 2.2.0) pipelines, which automatically and equally applied algorithms for primary object identification and subsequent pixel intensity quantification within identified primary objects to all evaluated pictures according to pre-defined parameters. After image quantification, CellProfiler pipelines were further used to automatically and equally apply background subtraction and color brightness adjustment to all pictures for respective qualitative display.

### Data processing and statistical analyses

Numerical raw data were processed as described in the respective sections with Microsoft Excel (© Microsoft, Redmond, USA; Office version 16.16.27). Dot plots were produced in Microsoft Excel, while box plots were created using the web-tool of BoxPlotR (http://shiny.chemgrid.org/boxplotr/). For box plots, center lines show the medians, box limits indicate the 25th and 75th percentiles as determined by R software, whiskers extend 1.5 times the interquartile range from the 25th and 75th percentiles, outliers are represented by dots and data points are plotted as open circles. If indicated, statistical analyses were conducted in R. First, to determine whether variables met the assumptions of linear models, Kolmogorov-Smirnov’s or Shapiro Wilk’s and Levene’s test or F-test for normal distribution and homogeneity of variance were performed, respectively. If the variables analyzed were normally distributed, Welch two sample t-test and independent one-way ANOVA were conducted in order to compare two and more than two groups, respectively. ANOVA was followed-up by pairwise comparisons using the Tukey-honest significant difference and Games-Howell tests for homogenous and non-homogeneous variances, respectively. In case analyzed variables were not normally distributed, the non-parametric equivalents, namely the Wilcoxon rank sum and Kruskal Wallis tests, were calculated. The Kruskal Wallis test was followed-up by multiple comparisons using the Wilcoxon rank sum test. Significance level was set at *p* < 0.05 and adjusted by default when multiple comparisons were performed. Significance levels are indicated in figures as follows: *: *p* < 0.05; **: *p* < 0.01, ***: *p* < 0.001, ****: *p* < 0.0001. Figures were assembled using Inkscape (© Free Software Foundation Inc., Boston, USA; version 0.92.2).

## Results

### Human iNSCs can be derived from blood cells of donors aged 0 to 101 years

Our previous study had provided first evidence that iNSCs generated from EPCs of 31 to 62-year-old donors exhibit a remarkable reduction of epigenetic age (Sheng et al., 2018). To study the reset of aging signatures in more detail, we first extended the donor cohort to 15 individuals covering a spectrum from 0 to 101 years of age (Supplementary Table S1). For comparison we generated from the same EPC samples isogenic iPSC-derived NSCs (iPS-NSCs; (Reinhardt et al., 2013), which – as iPSC derivative – undergo a complete age reset (Lo Sardo et al., 2017; Sheng et al., 2018). In both protocols, proliferation is sustained by promotion of sonic hedgehog and Wnt signaling, yielding NSCs with highly similar transcriptome and differentiation potential (Fig. 1A; Sheng et al., 2018). For iNSC generation, we employed a Sendai virus (SeV) system to achieve robust overexpression of SOX2 and cMYC. Following SeV transduction, EPCs downregulated the blood cell marker CD71 within few days, and expression of the NSC marker DACH1 could be detected in converting cells as early as 12 days post infection (Figure 1 B). Already at this time point, cells also expressed PAX6 (Figure 1 C). We verified that iNSCs and their isogenic iPS-NSC lines homogenously expressed the NSC markers SOX2 and NES even after transgene removal and extended *in vitro* cultivation (Figure 1 D; Supplementary Figure S1 A). Quality control by SNP analysis verified that iNSCs and iPS-NSCs remained genomically intact after cell programming and subsequent expansion for at least 20 passages, equaling to up to 6 months of continuous *in vitro* cultivation (Supplementary Figure S1 B). Furthermore, iNSCs and iPS-NSCs were successfully differentiated into TUBB3-positive neurons, S100β- and GFAP-positive astrocytes and O4-positive oligodendrocytes (Supplementary Figure S1 C).

**Figure 1:**
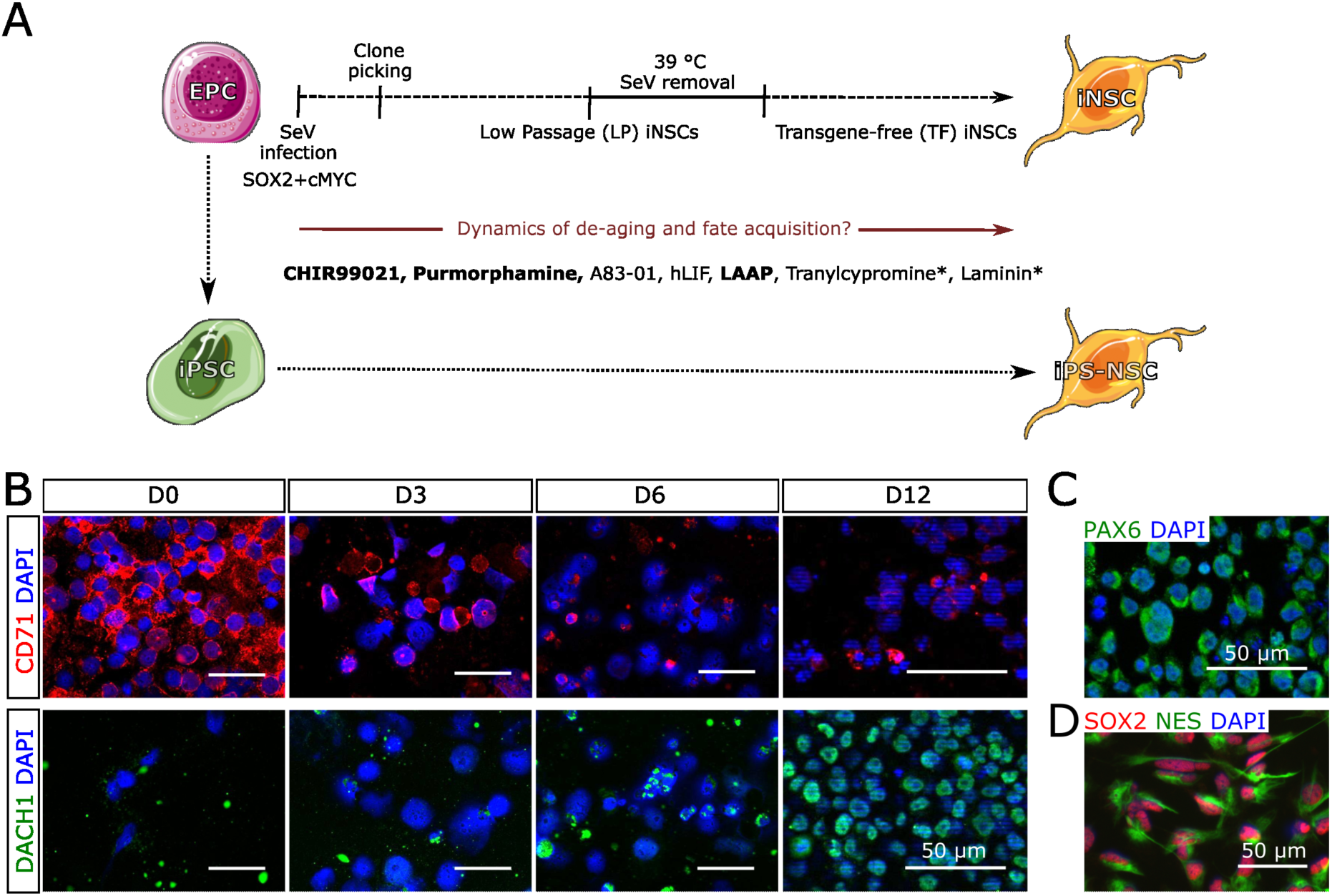
CD71-positive EPCs can be programmed into DACH1-expressing NSCs within less than two weeks. **(A)** Peripheral blood-derived EPCs were programmed into isogenic sets of directly converted iNSCs and iPS-NSCs. Samples were collected at different stages of iNSC conversion, for example around the time point of clone picking (D14), at low passage (LP) right before high temperature treatment and at a stage when established iNSC lines were transgene-free (TF), in order to investigate the dynamics of cellular age reprogramming in the process of iNSC conversion, using embryonic stage-like iPS-NSCs as controls. In the centrum of the figure, the components of the utilized iNSC media are listed. Components printed in bold are also constituents of the iPS-NSC medium. Components marked with asterisks are only used until day 10 of iNSC conversion. **(B)** Exemplary immunofluorescence images of cells stained against the EPC marker CD71 and the NSC marker DACH1 at different time points of iNSC conversion. Days indicate the time after SeV infection. Scale bars = 50 µm. Differences in cell size are most likely due to the cyto-spin procedure used to adhere suspension cells prior to the staining procedure. **(C)** At day 12 of conversion, arising iNSCs were stained against the NSC marker PAX6. Scale bar = 50 µm. **(D)** Representative immunofluorescence image of an established iNSC line stained against the NSC markers SOX2 and NES. This picture represents an excerpt of Supplementary Figure 1 A. Scale bar = 50 µm.

### iNSC conversion is associated with a protracted loss of aging signatures

We first assessed the epigenetic age of our enlarged cohort including source EPCs, low passage iNSCs and iPS-NSCs using the DNAm age prediction algorithm published by Horvath *et al* (2018). As expected, the DNAm age of EPCs highly correlated with the chronological age of the respective donors, demonstrating the accuracy of the age predictor algorithm. Concordant with published data, all iPS-NSC lines exhibited an embryonic-like epigenetic age, irrespective of their original donor age. Interestingly, iNSCs, too, exhibited a substantial degree of epigenetic de-aging across all donor ages, maintaining only around 13 % of the original donor signatures already at low passages (Figure 2 A). Intrigued by the reproducibility of this finding using donors of different ages, we next studied how age-associated signatures are affected by blood-to-iNSC transdifferentiation across time. Assessing the DNAm age dynamic in three iNSC lines derived from donors aged 50 to 101 years of age, we found that this de-aging represents a largely protracted process continuing for several weeks, and that it takes about 50 days of iNSC conversion until the majority of age-related epigenetic signatures are reset (Figure 2 B).

**Figure 2:**
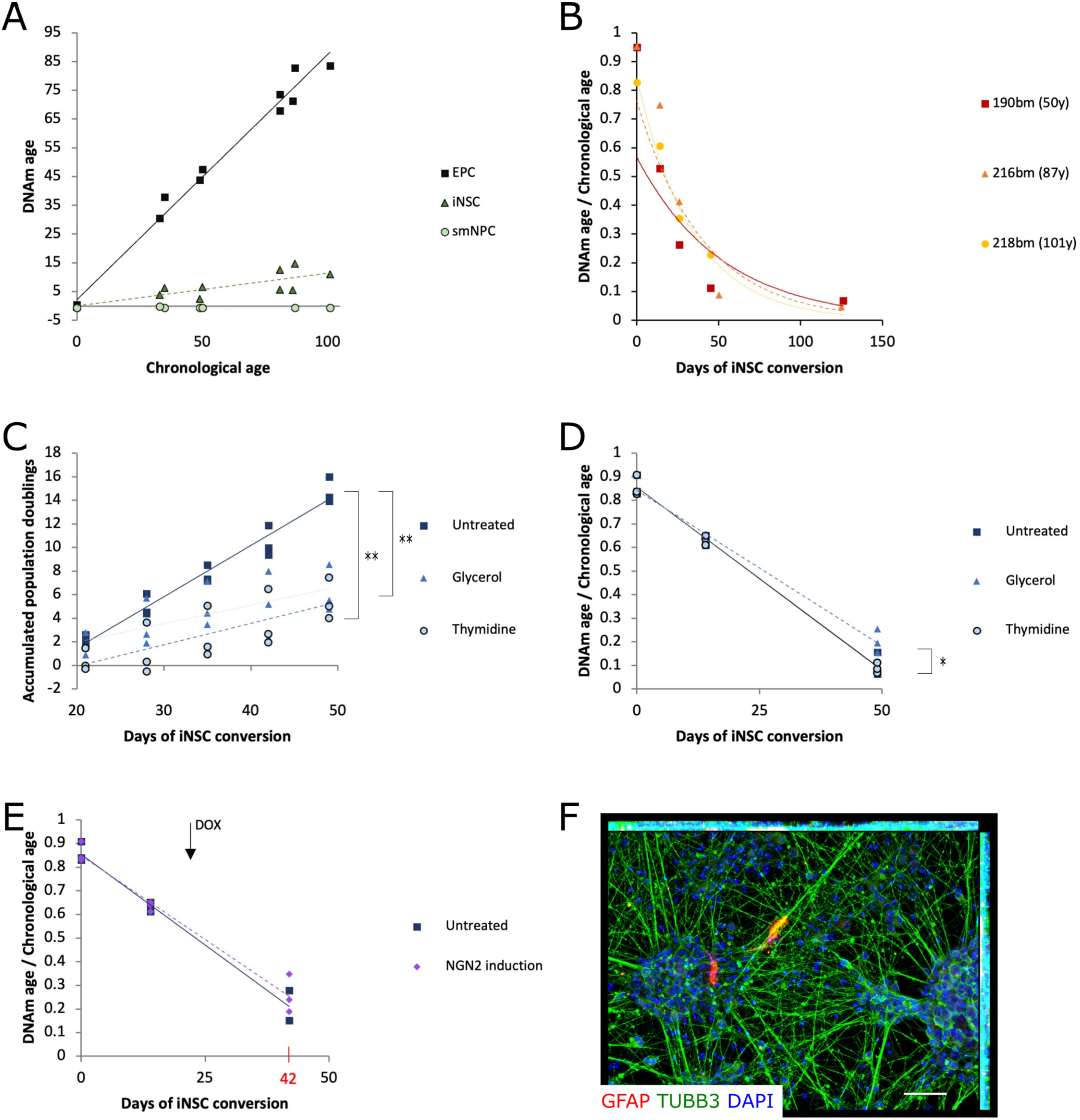
DNAm age dynamics upon iNSC conversion reveal that epigenetic de-aging is a protracted process that is largely unaltered by the manipulation of stem cell proliferation. **(A)** DNAm age predictions based on Horvath’s algorithm (Horvath et al., 2018) for EPCs and low passage blood-derived iNSCs as well as iPS-NSCs of donors of different ages. **(B)** Time course analysis of DNAm ages upon blood-to-iNSC conversion using three genetically distinct 50-101 years-old donor-derived cell preparations. **(C)** Graph depicting accumulated population doublings over the 5-week-long period of proliferation inhibition using continuous glycerol or cyclic thymidine treatment. N = 3 donors. Results of the Tukey ANOVA post-hoc test: Untreated vs. glycerol: *p* = 0.002; untreated vs. thymidine: *p* = 0.001 and glycerol vs. thymidine: *p* = 0.842. **(D)** Dot plot with trend lines depicting DNAm ages based on Horvath’s algorithm (Horvath et al., 2018), normalized to the chronological age of the respective donor, for untreated, glycerol-treated and thymidine-treated iNSCs. N = 3 donors. Results of the Tukey ANOVA post-hoc test: Untreated vs. glycerol: *p* = 0.056; untreated vs. thymidine: *p* = 0.977 and glycerol vs. thymidine: *p* = 0.044. **(E)** Dot plot with trend lines showing DNAm ages, normalized to the chronological age of the respective donor, after three-week-long NGN2-driven differentiation as compared to proliferating control cultures of early stage iNSCs. The arrow marks the day on which treatment with doxycycline (DOX), *i.e.,* induction of NGN2, started. N = 3 donors. Result of the Welch two sample t-test: *p* = 0.635. **(F)** Representative image showing that cultures mostly consisted of TUBB3-positive neurons and few GFAP-positive astrocytes after NGN2-mediated differentiation. N = 3 donors. Scale bar = 50 µm.

We confirmed the absence of cellular aging hallmarks in established, transgene-free (TF) iNSCs from donors of different ages (0, 87 and 101 years of age) by comparing them to their isogenic fully rejuvenated iPS-NSCs in different assays. First, we found that the nuclear lamina-associated *LMNB* and *LAP2α*, which are both negatively correlated with aging (Scaffidi & Misteli, 2006; Shah et al., 2013), are similarly expressed in both cell types and age groups. Expression levels of the age-associated genes *RANBP17*, *LAMNA* and *PCDH10* (Mertens et al., 2015), as well as the senescence-related genes *CDKN1a* and *CDKN2a* were similar in young and old donor-derived iNSCs and iPS-NSCs, too (Supplementary Figure 2 A). Although qualitative changes in LMNA/C expression upon aging are much better characterized than absolute differences in expression levels, expression of LMNA/C tended to be higher in iNSCs as compared to isogenic iPS-NSCs and in young as compared to old donor-derived cells – a trend we observed on both RNA and protein level (Supplementary Figure 2 A, B). Second, Western blot analysis of p62 abundance and LC3 conversion revealed no impairments of autophagy in iNSCs (Supplementary Figure 2 C). Finally, DNA damage and mitochondrial ROS production in iNSCs and iPS-NSCs were overall comparable (Supplementary Figure 2 D, E). Altogether, these data indicate that even iNSCs derived from very old donors are epigenetically and cellularly de-aged after direct conversion, highly resembling embryonic stage-like iPS-NSCs.

### Epigenetic de-aging upon iNSC conversion is not dependent on stem cell proliferation

Considering that epigenetic rejuvenation is observed upon reprogramming of somatic cells into self-renewing iPSCs (Lo Sardo et al., 2017) and direct conversion into proliferating iNSCs but not upon transdifferentiation into post-mitotic neurons (Huh et al., 2016), we hypothesized that de-aging might be driven by the persistent mitotic activity of stem cells. We thus performed two sets of experiments manipulating iNSC proliferation using cells derived from three aged donors (81-86 years of age). First, we treated iNSCs with glycerol – which is known to reduce proliferation of various cell types – or thymidine – which is commonly used as a cell cycle-synchronizing agent, since it arrests cells in the G1/S phase of the cell cycle – starting from day 14 of conversion. Despite their different modes of action, both treatments resulted in significantly reduced accumulated population doublings at day 49 of conversion (Figure 2 C). Yet, neither the relative DNAm age of glycerol-nor thymidine-treated cultures differed significantly from untreated cultures, although there was a slight but statistically significant increase in the relative DNAm age of glycerol-treated cultures as compared to thymidine-treated cultures (Figure 2 D).

Second, we infected EPC-derived iNSCs at day 7 of conversion with lentiviruses encoding a doxycycline-inducible NGN2 expression system. Forced induction of NGN2 has been shown to rapidly induce neuronal differentiation of human NSCs (Ho et al., 2016). After 14 additional days of expansion, a fraction of the infected cells was kept in proliferation, while the rest was subjected to DOX induction under differentiation-promoting conditions. At day 42 of conversion (*i.e.*, after 20 days of differentiation), the relative DNAm age of NGN2-overexpressing, differentiated cultures did not significantly differ from proliferative iNSCs (Figure 2 E), despite the fact that cultures forced to differentiate by NGN2 overexpression indeed mostly consisted of TUBB3-positive neurons and few GFAP-positive astrocytes (Figure 2 F). In sum, our data thus suggest that epigenetic de-aging upon iNSC conversion is protracted, and does not depend on sustained cell proliferation, even extending into a post-mitotic stage.

### Protracted consolidation of iNSC identity

We next assessed DNAm changes in the process of iNSC conversion at a larger scale, including data from the cell lines programmed in the course of this project as well as data produced and published in the course of our previous publication (Sheng et al., 2018). In total, we included DNAm data from EPCs of twelve donors as well as from the low passage (LP; P3-P5) iNSC lines established thereof. For a subset of these donors, we additionally included samples collected in the early phase of iNSC conversion, *i.e.*, from emerging neuroepithelial-like cultures at days 14 (N=6) and 26 (N=4) of conversion, as well as transgene-free iNSCs after extended cultivation (TF; P21; N=3). After raw data filtering, in total 728,674 probes were compared across these 37 samples. We found that while EPCs and day 14-iNSCs form very distinct clusters in the multidimensional scaling plot (Figure 3 A), samples collected at later time points appear quite similar to each other. We then mapped significant most variable positions (MVPs; Benjamin Hochberg-adjusted p<0.05) to genes, assessing how many were commonly affected by the analyzed contrasts across time in conversion (Figure 3 B). Subsequent Reactome pathway analysis revealed a broad spectrum of pathways, of which most could be assigned to one of five categories: (i) neuronal system, especially neuronal migration, neurite outgrowth and pathfinding; (ii) cell migration, adhesion and division in multiple tissues, including pathways relevant to extracellular matrix (ECM) and mesenchymal-to-epithelial transition (MET); (iii) signal transduction, (iv) angiogenesis and (v) the hematopoietic system (Supplementary Figure 3). Except for signal transduction and the hematopoietic system, terms of the other three categories were found to be regulated across all stages of iNSC conversion. We then conducted gene set enrichment analysis directly on the identified differentially methylated regions (DMRs; FDR< 0.05). Interestingly, focusing on the 45 gene ontology (GO) terms that were commonly enriched across all stages of iNSC conversion and *in vitro* cultivation, 14 were associated with nervous system development such as ‘neurogenesis’, ‘axon development’ and ‘neuron projection morphogenesis’ (Supplementary Table S2). In this analysis, morphogenesis-, cytoskeleton- and signal transduction-associated terms accounted for the majority of the remaining commonly regulated GO terms as well.

**Figure 3:**
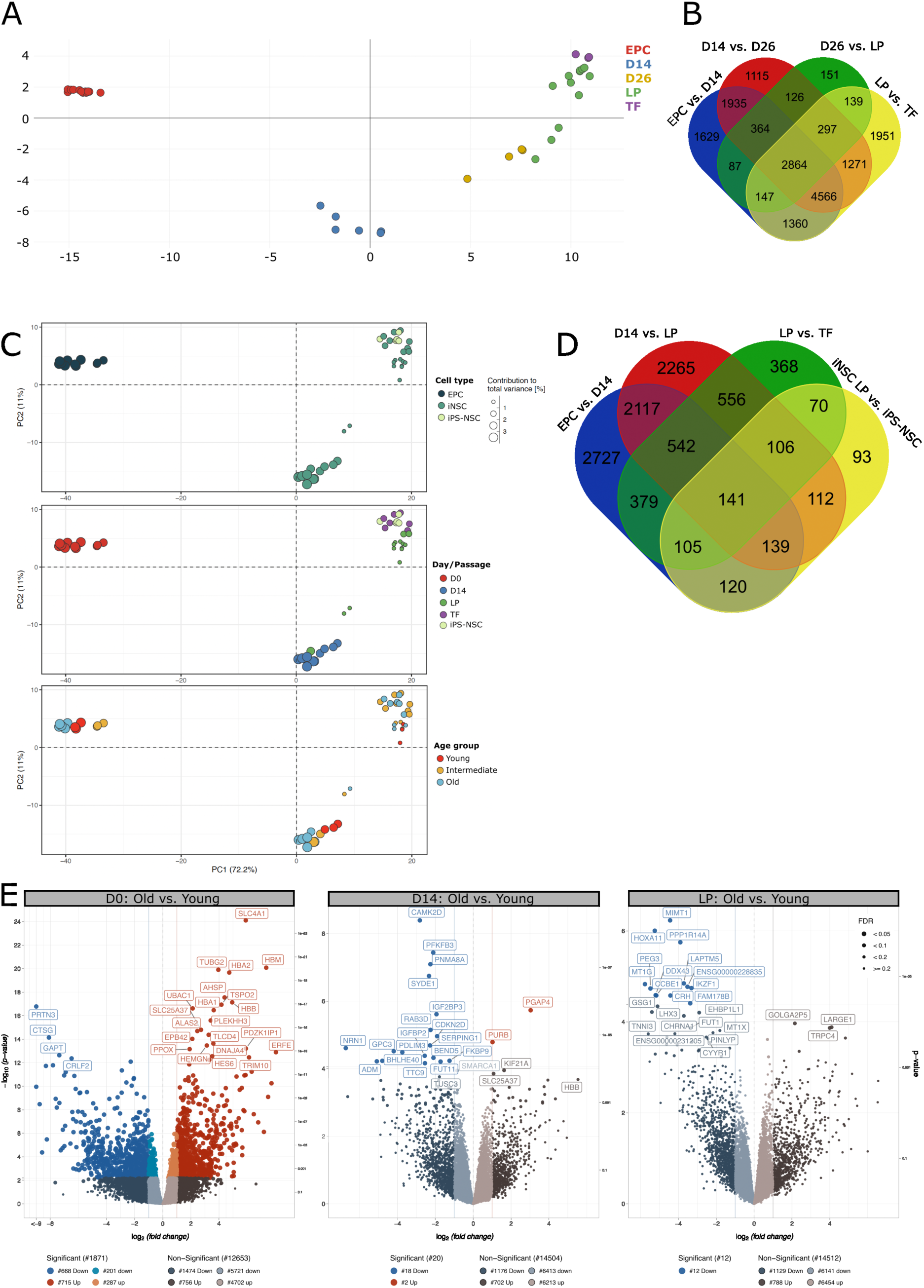
Epigenetic and transcriptomic consolidation of NSC identity is a protracted process, which extends at least beyond day 26 of conversion. **(A)** DNAm data-derived multidimensionality scaling plot based on the top 1,000 most variable positions (MVPs) across samples. **(B)** Venn diagrams depicting the number of genes, which the significant MVPs were mapped to, for all performed contrasts. iNSCs at LP: P3-P5; iNSC at TF: P21. **(C)** RNA sequencing data-derived principal component analysis (PCA) plots color-coded according to each sample’s cell type (upper PCA), the stage (*i.e.*, day of conversion or passage) at which the respective sample was harvested (middle PCA) or the age assignment of the original donor of each sample (lower PCA). **(D)** Venn diagram showing the number of differentially expressed genes (DEGs) identified in relevant contrasts across cell types (*i.e.*, iPS-NSC vs. iNSC, both at LP) or stages of conversion (*i.e.*, day 0 vs. 14 of conversion, day 14 vs. LP and LP vs. TF). iNSCs at LP: P4-P6; iNSC at TF: P18-P21; iPS-NSC: P6. **(E)** Volcano plots representing DEGs comparing young to old donor-derived cells at a specific stage of iNSC conversion (*i.e.*, in the original blood cells at day 0 of conversion, at day 14 of conversion and in LP iNSCs). Categorization according to donor age: young: 0 years of age; intermediate: 31-50 years of age; old: 62-101 years of age.

In analogy to the DNAm analyses, we also assessed global transcriptomic changes upon iNSC conversion. Specifically, we included twelve EPC samples as well as ten day 14, fourteen LP (P4-P6) and seven TF (P18-P21) iNSC samples. For five donors, we additionally included isogenic iPS-NSC samples at LP (P6) as reference. Principal component analysis (PCA) of this set of RNA sequencing dataset revealed that while all EPC samples formed one distinct cluster, iNSC samples were distributed across two main clusters. One cluster predominantly consisted of day 14 iNSCs, whereas the other cluster was composed of LP and TF iNSCs, with the latter intermingling with iPS-NSCs (Figure 3 C). Specifically, while 6,270 and 5,978 genes were altered in the early phase of iNSC conversion comparing EPCs to day 14 iNSCs and day 14 to LP iNSCs, respectively, only 2,267 genes were differentially expressed comparing LP to TF iNSCs (Figure 3 D).

Notably, when assigning all donors to young (0 years of age), intermediate (31-50 years of age) or old (62-101 years of age) donor age groups, we noticed already in the PCA that while young, intermediate and old-age-donors slightly segregated at the EPC stage, they progressively intermingled at later stages of iNSC conversion (Figure 3 C). Retrieving DEGs between age groups per analyzed time point of iNSC conversion, we found that the expression of 1,871 genes significantly differed in starting blood cells derived from young and old donors at day 0 of conversion. However, only 20 and 12 genes were differentially expressed between age groups at day 14 of conversion and LP, respectively (Figure 3 E). This observation suggests that cells derived from donors of different ages become transcriptomically more similar to each other over time in conversion.

Since the main transcriptomic shift along iNSC conversion could thus not be explained by the observed de-aging phenomenon, we finally performed GO term enrichment by overrepresentation analysis on the genes that were altered between different stages of iNSC conversion. This analysis revealed that genes that were up- and downregulated during the first 14 days of iNSC conversion were almost exclusively associated with the neural lineage and DNA replication and repair, respectively (Figure 4 A). Afterwards, from day 14 to LP, genes associated with translation were downregulated, which could suggest that the majority of proteomic changes relating to the blood-to-iNSC fate shift are indeed achieved before LP. For the upregulated genes, 6 out of the top 10 GO terms related to the nervous system (Figure 4 B). From LP to the TF stage, genes associated with viral response were downregulated next to translation, which is compatible with the removal of SeV transgenes by high temperature treatment occurring during this period. On the other hand, 5 of the top 10 GO terms relating to upregulated genes were still directly or indirectly associated with the nervous system (Figure 4 C).

**Figure 4:**
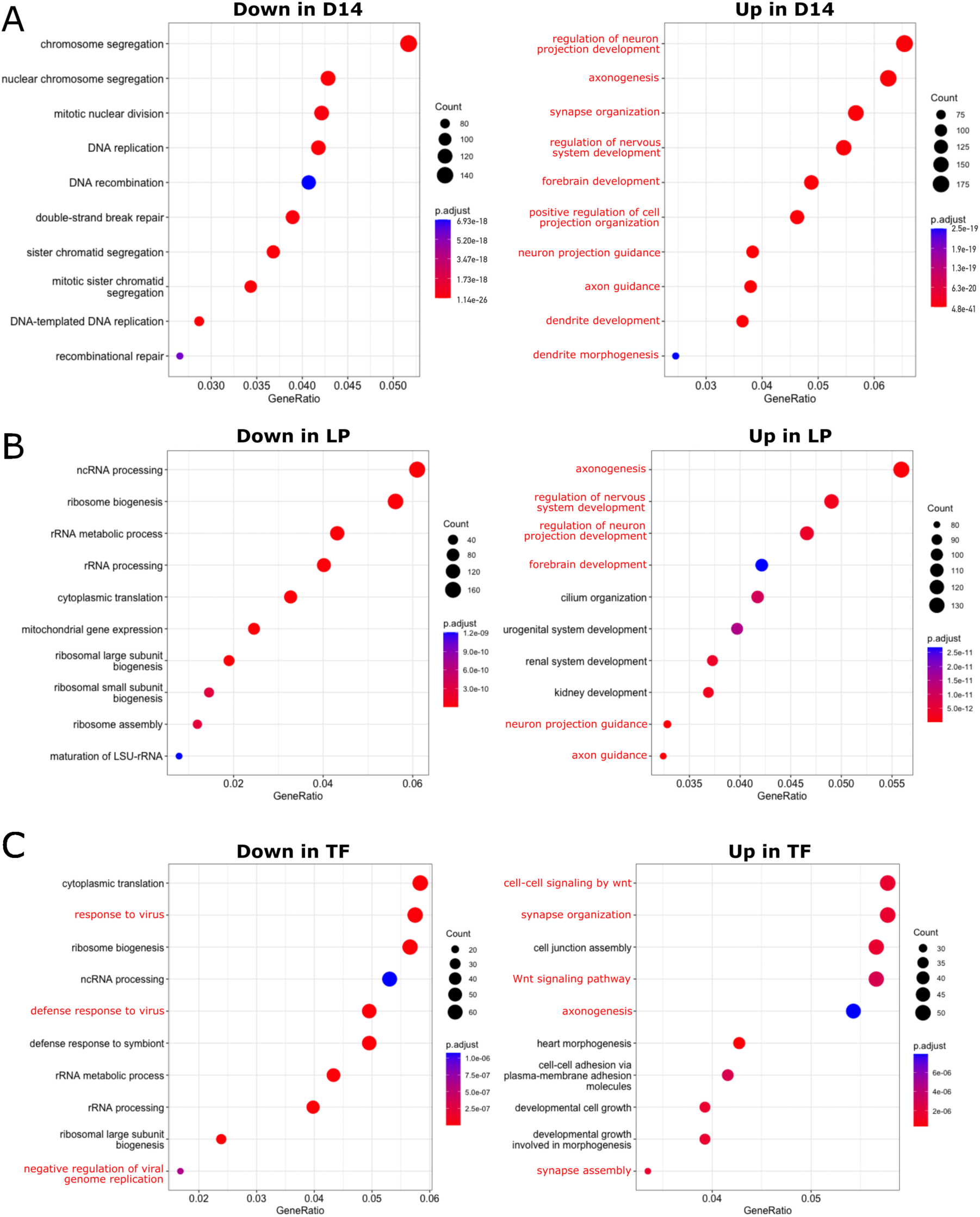
Blood-to-NSC transdifferentiation successively induces transcriptomic changes associated with the neural lineage, as well as cell cycle and translation. Overrepresentation analyses were performed for significant GO terms relating to DEGs from the day 0 vs. day 14 (A), day 14 vs. LP (B) and LP vs. TF (C) contrasts. The top 10 up- and down-regulated terms are depicted per contrast. Neuronal system-associated GO terms are printed in red.

In sum, these data indicate that despite the early acquisition of an NSC-characteristic morphology and expression of characteristic NSC markers within days after SeV infection, the full transcriptomic and epigenetic consolidation of cellular identity takes weeks after initiation of the blood-to-NSC fate shift.

## Discussion

### Profound and protracted de-aging extending up into the postmitotic stage

The results of our study show that direct conversion of peripheral blood-derived EPCs into iNSCs is associated with pronounced epigenetic de-aging. iNSCs generated via temporary overexpression of SOX2 and cMYC retain less than 5 % of the DNAm age of their original blood cells, which were obtained from donors aged 0 to 101 years of age. Unexpectedly, this de-aging process is largely protracted, stretching across the first 50 days of iNSC conversion.

Epigenetic de-aging has also been observed during classic iPSC reprogramming, reaching a complete reset within approximately twenty days (Gill et al., 2022; Olova, Simpson, Marioni, & Chandra, 2019). However, in the case of iNSC conversion, this process follows much slower dynamics, reaching a de-aging level down to about 20 % of the chronological age only after around 7 weeks. Once established as expandable cell populations, iNSCs generated even from very old donors (87-101 years of age) are devoid of classic age-associated cellular changes such as alterations of the nuclear lamina, impairments of the autophagic flux and mitochondrial function, as well as hallmarks of senescence. All of these properties were comparable to isogenic iPS-NSC, which, due to their iPSC origin, can be considered cellularly rejuvenated. Taken together, these data corroborate our conclusion that iNSC conversion yields cells devoid of aging signatures.

Interestingly, epigenetic de-aging in the context of direct transcription factor-based neural conversion seems not be restricted to the SOX2/cMYC scenario reported here. Scrutinizing the literature on this topic, we came across the work of Thier *et al*. (2019), who used BRN2, KLF4, ZIC3 and SOX2 to convert human fibroblasts into neural plate border stem cells (NPSCs). When we ran two different algorithms for DNAm age calculation (Horvath, 2013 and Horvath et al., 2018) on their publicly available DNAm data, we found that the human source fibroblasts predicted to be mid-age (46-48 and 33-38 years, respectively) yielded vastly de-aged NBSCs (1-2 and 0.4-0.8 years, respectively; data not shown). Hence, it seems that not only blood-to-iNSC but also human fibroblast-to-NPSC conversion is associated with an epigenetic de-aging process. Considering that overexpression of SOX2 is the only commonality of both conversion paradigms, this transcription factor might merit further investigation in the context of somatic cell de-aging. Notably, though, other studies have linked expression of Myc to cellular aging, too (Neumann et al., 2021).

Remarkably, when newly generated iNSCs were forced to differentiate into neurons, the epigenetic de-aging process continued even in the postmitotic stage. To us this observation came as a surprise, since reprogramming into iPSCs and conversion into iNSCs are both associated with proliferation, whereas direct transcription factor-based conversion of fibroblasts into iNs without an intermediate proliferation step results in epigenetic age preservation (Huh et al., 2016). The continuation of epigenetic de-aging into the postmitotic neuronal stage could further suggest that this process is independent of cell fate conversion. In that regard, our transdifferentiation system might serve as a blueprint for future studies focusing on potential drivers of somatic cell rejuvenation.

### Prolonged consolidation of iNSC DNAm and transcriptome signatures

When we looked at the quality of the DNAm changes in the course of conversion, we found that most regulated pathways were either associated with classifications relating to nervous system, blood, cell morphogenesis and cytoskeleton or cell function, including migration, adhesion, division and signaling. Considering that the conversion process comprises a transition from a mesenchymal to an epithelial-like cell population where freely moving blood progenitor cells are transdifferentiated into an adhesive neuroepithelial cell population, changes in classifications such as cell morphology, adhesion and migration are to be expected. However, here we were again surprised to see that the changes continued across almost all stages of iNSC conversion, up to the stage when iNSCs are already transgene-free and long-term expanded.

At the transcriptomic level, too, changes continued for an extended period of time. Signatures indicating an early neural fate entry were largely established in the first two weeks, when newly generated iNSCs had reached a stably proliferating PAX6- and DACH1-positive fate. However, categories regarded as prerequisite for further neuronal maturation such as axonogenesis and synapse organization continued to go up until the very late transgene-free stages – despite the fact that cells were in proliferating conditions at all times. Other DEG clusters such as, for example, Wnt signaling might in part be due to the specific culture conditions promoting these pathways. This is also true for viral response signatures at the transition from LP to TF stage, which are compatible with SeV removal.

Taken together, our data point to an unexpectedly protracted fate acquisition and consolidation process in EPC-to-NSC programming, which is accompanied by prolonged epigenetic de-aging.

In this context, it is worth noting that PCA of our RNA sequencing data suggests that transcriptome-based separation of young, intermediate and old donors is only possible before initiation of SOX2/cMYC overexpression. Although our analysis is limited by comparably small group sizes and we cannot rule out that the observed fate shift-associated changes, which were much stronger, may overshadow de-aging-associated changes, this observation could suggest that transcriptome de-aging might be achieved faster than the loss of epigenetic signatures and that transcriptomic changes are primarily induced by the overexpressed transcription factors as such rather than epigenetic changes. In a recent study on pan-mammalian and pan-tissue epigenetic aging markers, the GO term ‘nervous system development’ was found to be highly significant for CpG sites that are positively correlated with aging (Lu et al., 2023). In addition, positively as well as negatively age-related CpG sites were enriched in targets of SOX2, MYC and OCT4 (Lu et al., 2023). This association merits careful further investigation, for instance with respect to the effects of transgene repression on cellular identity and age at different stages of neural conversion.

### SOX2/cMYC-induced transdifferentiation and de-aging: Uncoupled processes or evidence of a pre-pluripotent stage?

In a broader context, it is worth considering that both SOX2 and cMYC are important components of iPSC reprogramming, and ‘partial’ or ‘interrupted’ reprogramming has been employed in numerous *in vitro* and *in vivo* settings to induce cellular rejuvenation and organismal regeneration (for review see Puri & Wagner, 2023; Simpson, Olova, & Chandra, 2021; Singh & Zhakupova, 2022). Interestingly, it was lately reported that iNSCs converted from tail tip fibroblasts of 30 month-old mice show a highly variable degree of DNAm age reset. After overexpression of an engineered Sox17 (eSox17^FNV^) variant in conjunction with Klf4 ± Brn2 and cMyc, which was shown to converge on the same mechanism as conversion with Sox2, namely dimerization with Brn2 on the canonical SoxOct motif, fibroblast-derived iNSCs attained a DNAm age ranging from 6.9 to 27.9 months (Weng et al., 2023). Thus, while some clonal lines seem to have retained the majority of the age-related DNAm signatures of their source cell, others underwent epigenetic de-aging in the process of conversion. Intriguingly, the authors propose that the addition of cMyc to the two-factor-cocktail Klf4 plus eSox17 ^FNV^ might promote the acquisition of an Oct4-positive, Nanog-negative pre-pluripotent state (Weng et al., 2023). However, whether or not the transition through a pre-pluripotent stage was associated with or even necessary for de-aging was not investigated in this study.

The results of our previous study provided no evidence of a pluripotency transit of SOX2/cMYC-induced iNSCs (Sheng et al., 2018). Yet, the emergence of kidney, urogenital and heart signatures detected in our current RNA sequencing data could be compatible with the concept partial reprogramming, where the media conditions eventually lead the cells into a neural fate. Notably, though, these signatures were intermediately regulated at low passage, *i.e.,* not within the first weeks of conversion, when one would expect a potential pluripotency transit to occur. Furthermore, emergence of aberrant fates has also been observed in the context of reprogramming to and transcription factor-mediated differentiation of iPSCs, as well as direct fibroblast-to-iN conversion (Deng et al., 2021; Lin et al., 2021; Treutlein et al., 2016). Thus, while the available data argue against a pluripotent transit in the context of iNSC programming, extensive fine-grained time course studies with single cell resolution will be required to determine whether and how close emerging iNSCs approximate pluripotency and to what extent such an approximation is required for the observed de-aging phenomenon.

## Conclusion

In summary, our study reveals that both, consolidation of a NSC fate as well as (epigenetic) de-aging are protracted processes in transcription factor-based conversion of human blood cells. This prolonged time window of several weeks may provide a unique opportunity for mechanistic studies that aim at disentangling both processes and delineating the drivers for somatic rejuvenation. In that regard, our conversion system might serve as an alternative to partial or interrupted iPSC reprogramming and its exploration for cellular de-aging.

## Acknowledgements

We specifically acknowledge Cornelia Thiele and Melanie Bloschies (all Institute of Reconstructive Neurobiology, University of Bonn) for outstanding technical support. We further thank Ullrich Wüllner (Clinic and Polyclinic for Neurology, University Hospital Bonn), Anja Schneider (Clinic for Neurodegenerative Diseases and Gerontopsychiatry, University Hospital Bonn) and Waltraut Merz (Department of Obstetrics and Prenatal Medicine, University Hospital Bonn) for aid during subject recruitment and blood withdrawal. Lentiviruses and lentiviral plasmids for rtTA TRE-regulated NGN2 were kindly provided by Laura Stappert (formerly Institute of Reconstructive Neurobiology, University of Bonn). We would like to thank the Next Generation Sequencing Core Facility and the Core Unit for Bioinformatics Data Analysis of the Medical Faculty at the University of Bonn for providing support and instrumentation, funded by the Deutsche Forschungsgemeinschaft (DFG, German Research Foundation). Furthermore, many thanks to the Neuropathology Department of the University Hospital Bonn, and especially Alexandra Brüggemann, for helping with the centrifugation of suspension cells for staining. Finally, we acknowledge Michael Ziller (Max Planck Institute of Psychiatry, Munich) for insightful discussions in the course of this project.

This work was supported by the European Union’s Horizon 2020 research and innovation program (grant agreement no. 874758 NSC-Reconstruct to O.B.), the DFG (grants CRU344/417911533 and SFB 1506/1 to W.W.) and the German Federal Ministry of Education and Research (grant 01EK1603A-Neuro2D3 to O.B.; VIP+ PluripotencyScreen to W.W.).

## Authors’ contributions

L.J.B., C.S., M.P., A.T., W.W. and O.B. conceived and designed the study. L.J.B., R.K. and C.S. performed cell culture experiments. L.J.B., J.F. and A.B. analyzed and assembled data. L.J.B., A.B., M.P., A.T., W.W. and O.B interpreted data. L.J.B. and O.B and wrote the manuscript. All authors read and approved the final manuscript version.

## Data availability statement

The accession number of the DNA methylation data set published in Thier *et al*. (2019) is E-MTAB-5808, available via ArrayExpress. The accession numbers for the data reported in this paper are GSE295849 (DNA methylation data, available via GEO after journal publication) and phs004047.v1.p1 (RNA sequencing data, available via dbGaP after journal publication).

