## Supplemental Material for "Protracted fate acquisition and epigenetic de-aging during induced neural stem cell conversion of human blood cells"

### Supplementary Information

**Supplementary Methods**

***NSC differentiation***

In order to differentiate NSCs spontaneously, cells were plated in their respective cultivation medium on Matrigel-coated cell culture formats (1:30-1:45 dilution in DMEM/F12) at a density of $8.3 \times 10^4$ cells/cm$^2$. The day after NSC plating, medium was changed to NGMC (1:1 DMEM/F12 : Neurobasal with 0.5x N2 supplement, 0.5x B27 supplement without vitamin A, 1 mM L-glutamine, 800 µg/ml D(+)glucose (Carl Roth), 1x Pen/Strep and 0.5 mM dbcAMP (Sigma-Aldrich)) containing 10 ng/ml BDNF, 10 ng/ml GDNF (both Cell Guidance Systems) and 64 µg/ml LAAP. DAPT (Axon Biotech) was added to the differentiation medium at a concentration of 5 µM up until days 10-14 of differentiation in order to promote cell cycle exit. Medium was changed three times a week during the 6-week-long differentiation period.

To generate oligodendrocytes, a multi-stage differentiation paradigm published by (Gorris et al., 2015) was adapted. In short, cells were plated at a density of $8.9 \times 10^4$ cells/cm$^2$ onto Matrigel-coated cell culture-dishes (1:60 dilution in DMEM/F12). The day after, medium was changed to oligodendrocyte differentiation medium stage I (oligodendrocyte base medium consisting of DMEM/F12 plus 1x N2 supplement, 2 mM L-glutamine, 1.6 mg/ml D(+)glucose, 200 µg/ml Apo-transferrin and 20 µg/ml insulin (both Sigma-Aldrich), supplemented with 10 ng/ml EGF (Sigma-Aldrich), 10 ng/ml PDGF-AA (Bio-Techne), 10 µM forskolin (Tocris Bioscience) and 1 µM SAG (Merck Millipore, Burlington, USA)). On day 7 of oligodendrocyte differentiation, cultures were replated onto Matrigel-coated cell culture-dishes (1:30 dilution in DMEM/F12) at a density of $\sim 1 \times 10^4$ cells/cm$^2$. On day 14, medium was switched to oligodendrocyte differentiation medium stage II (oligodendrocyte base medium supplemented with 1x B27 supplement without vitamin A, 10 ng/ml PDGF-AA, 30 ng/ml T3 (Sigma-Aldrich), 200 ng/ml Noggin (Bio-Techne) and 200 µM ascorbic acid (Sigma-Aldrich)). One week later, medium was changed to oligodendrocyte differentiation medium stage III (oligodendrocyte base medium supplemented with 1x B27 supplement without vitamin A, 60 ng/ml T3, 200 µM ascorbic acid, 10 ng/ml rhIGF1, 1 µg/ml laminin and 10 ng/ml NT-3 (PeproTech Inc, Rocky Hill, USA)). Medium was changed three times a week during the in total 7-week-long differentiation period.

***SNP analysis***

DNA extraction was performed as described in the main manuscript text. For single nucleotide polymorphism (SNP) analysis, DNA samples were diluted to a concentration of 55 ng/µl with

AE buffer (Qiagen) and provided to the Next Generation Sequencing Core Facility of the University of Bonn. Samples were then processed by whole-genome amplification, fragmentation and subsequent DNA-oligomer-hybridization and SNP genotyping on Illumina's (San Diego, USA) Infinium Global Screening Array platform versions 1-3. Raw data were processed using the associated GenomeStudio software using chip version-specific parameters. Samples with a call rate below 0.9 were excluded from further analysis. All chromosomes, except the gender-specific X- and Y-chromosomes, were assessed for copy number variations (CNVs). All established cell lines were compared to their cell line of origin (*i.e.*, iPSCs and iNSCs were compared to their respective EPCs of origin, whereas iPS-NSCs were compared to the respective iPSCs of origin) in order to identify CNVs, which were introduced by the according intervention (*e.g.*, cell programming). Cell lines were considered genomically intact, if all newly introduced CNVs were estimated with a confidence score lower than 200 as provided by GenomeStudio and/or were smaller than 500 kb.

***Quantitative reverse transcription real-time polymerase chain reaction (qPCR)***

cDNA was synthesized from 1 µg RNA using Quanta Biosciences' (Beverly, USA) qScript kit according to the manufacturer's instructions. The concentration and purity of cDNA samples were analyzed with a Nanodrop 2000c. qPCRs were performed on an Eppendorf (Hamburg, Germany) Mastercycler ep*gradient* S realplex in 96-well-format with 300 ng cDNA per reaction. Primer sequences were as follows: *18S:* F-5'-TTCCTTGGACCGGCGCAAG-3', R-5'-GCCGCATCGCCGGTCGG-3'; *CDKN1a*: F-5'-TGACCCTGAAGTGAGCACAG-3', R-5'-AAGGTACAGGGGAGCCAAAG-3'; *CDKN2a-p16-INK4a*: F-5'-GGGTCGGGTAGAGGAGGTG-3', R-5'-ACCGTAACTATTCGGTGCGT-3'; *CDKN2a-p14-ARF*: F-5'-TCTTGGTGACCCTCCGGATT-3', R-5'-CGGGATGTGAACCACGAAAAC-3'; *LAMNA*: F-5'-CACCTGGAACTGGACACAGA-3', R-5'-AGGGACAGGGATGTGATTGA-3'; *LAP2a*: F-5'-GCAGGCAGACATTAGTCAAGC-3', R-5'-CGACCTACAGTGGCATTTCC-3'; *LMNA*: F-5'-GCTCTTCTGCCTCCAGTGTC-3', R-5'-ACATGATGCTGCAGTTCTGG-3'; *LMNB*: F-5'-GAGGTTGCTCAAAGAAGTACAGTC-3', R-5'-TTACATAATTGCACAGCTTCTATTG-3'; *LMNC*: F-5'-CTCAGTGACTGTGGTTGAGGA-3'; R-5'-AGTGCAGGCTCGGCCTC-3'; *PCDH10*: F-5'-ATGCCTTCTTTTGTCCCTTCT-3', R-5'-AGTCCATCCAGCTCCTTCC-3'; *RANBP17*: F-5'-GGATCCTGGATTGAGACGAA-3', R-5'-GTGCTTCCAGGCTCGTTCTA-3' (all from Thermo Fisher Scientific or IDT (Coralville, USA)). Ct values of the genes of interest were normalized to the house-keeping gene *18S* and transformed to mean fold changes using the equation $2^{-\Delta\Delta Ct}$ (according to (Livak & Schmittgen, 2001). QPCR products were size-validated on 1.5 % agarose (Peqlab Biotechnologie GmbH) gels containing ethidium bromide (Sigma-Aldrich; 1:10,000 dilution).

***Western blotting***

In order to assess autophagic flux, cells were first stimulated with 40 nM bafilomycin A (BAFA; Enzo Biochem Inc) for 4 hours at 37 °C. Afterwards, cells were washed once with 1x DPBS before being mechanically detached with a cell scraper and transferred to 1.5 ml-tubes in 1 ml ice-cold 1x DPBS. Samples were centrifuged for 5 minutes at 500 rcf and 4 °C. Supernatants were discarded and cell pellets were either directly processed or frozen in liquid nitrogen and stored at -80 °C.

For protein extraction, cell pellets were lysed in 45 µl RIPA buffer (25 mM Tris-HCl (pH 7.6), 150 mM NaCl, 0.1 % sodium dodecyl sulfate (SDS; all from Carl Roth), 1 % sodium deoxycholate (Sigma-Aldrich) and 1 % nonoxiol 40 (Thermo Fisher Scientific)) supplemented with 1x protease and phosphatase inhibitor cocktail (Thermo Fisher Scientific). Protein concentrations were determined with Roti-Quant reagent (Carl Roth). Absorbance at 595 nm (10 nm bandwidth, 25 flashes) was recorded using a Tecan (Männedorf, Switzerland) Infinite M Plex plate reader. A standard curve was fitted to the data points of BSA standards and used to calculate the protein concentration of all other samples.

For Western blotting, 20 µg protein, diluted in 4x Laemmli buffer (20 mM Tris-HCl, 10 mM EDTA, 10 mM acetic acid (Carl Roth), 20 % glycerol, 8 % SDS and 1 % bromophenol blue (Sigma-Aldrich)) supplemented with 10 % 2-mercapto ethanol (Sigma-Aldrich), were separated on home-made 15 % acryl amide SDS-Tris gels after denaturation at 95 °C for 5 minutes. Afterwards, samples were transferred to 0.2 µm polyvinylidene fluoride (PVDF) membranes (Bio-Rad Laborartories Inc). After blotting, membranes were washed once with 1x TBS-T (25 mM Tris-Base (pH 7.4) plus 150 mM NaCl (Carl Roth), supplemented with 0.1 % Tween-20 (Sigma-Aldrich)) before being stained with Amido-Black solution (0.2 % amido-black (Carl Roth), 10 % acetic acid and 40 % methanol) in order to visualize protein loading. After removing the Amido-Black stain with 2 % SDS, membranes were cut at the height of the 25 kDa marker (Page ruler pre-stained protein ladder; Thermo Fisher Scientific). The cut parts were separately incubated with 10 % milk powder (Carl Roth) in 1x TBS-T for 10 and 30 minutes for the low and high molecular weight parts, respectively, before primary antibodies diluted in 5 % milk powder in 1x TBS-T were incubated overnight at 4 °C: mouse monoclonal IgG to p62 (1:1,000; Abnova, Taipeh, Taiwan), mouse monoclonal IgG to LC3B (1:200; Enzo Biochem Inc) and mouse monoclonal IgG to GAPDH (1:1,000; Santa Cruz Biotechnology, Dallas USA). The next day, membranes were washed three times for 5 minutes with 1x TBS-T, before incubation with secondary antibodies diluted in 5 % milk powder in 1x TBS-T was performed for 1 hour at room temperature: HRP-linked anti-mouse IgG (1:7,500; Cell Signaling Technology, Danvers, USA). Three washing steps with 1x TBS-T and one final washing step with 1x TBS, each conducted for 5 minutes, followed secondary antibody incubation. Finally, membranes were subsequently developed using Classico (Merck Millipore) and Femto (Thermo Fisher Scientific) HRP substrates before documentation using a ChemiDoc XRS+ (Bio-Rad Laborartories Inc).

***Flow cytometry***

To quantify mitochondrial ROS production using MitoSOX staining, iPS-NSCs and iNSCs were seeded at a density of $5x10^4$ cells/$cm^2$. The next day, medium was changed to antioxidant-free NSC medium consisting of 1:1 DMEM/F12 : Neurobasal with 0.5x N2 supplement, 0.5x B27 supplement without antioxidants (Thermo Fisher Scientific), 2 mM L-glutamine, 0.0025% BSA, 3 µM CHIR99021 and 0.5 µM purmorphamine. After 2 days in antioxidant-free medium, the supernatant was discarded and cells were incubated with HBSS ± 5 µM MitoSOX Red (both Thermo Fisher Scientific) for 10 minutes at 37 °C. Afterwards, cells were washed once with 1x DPBS before being detached using ice-cold 0.5 mM EDTA. Fluorescence was measured using a FACSCalibur (BD Bioscience) or an Accuri C6 Plus flow cytometer (BD Bioscience). Gating was performed according to unstained negative control.

### Supplementary Figures

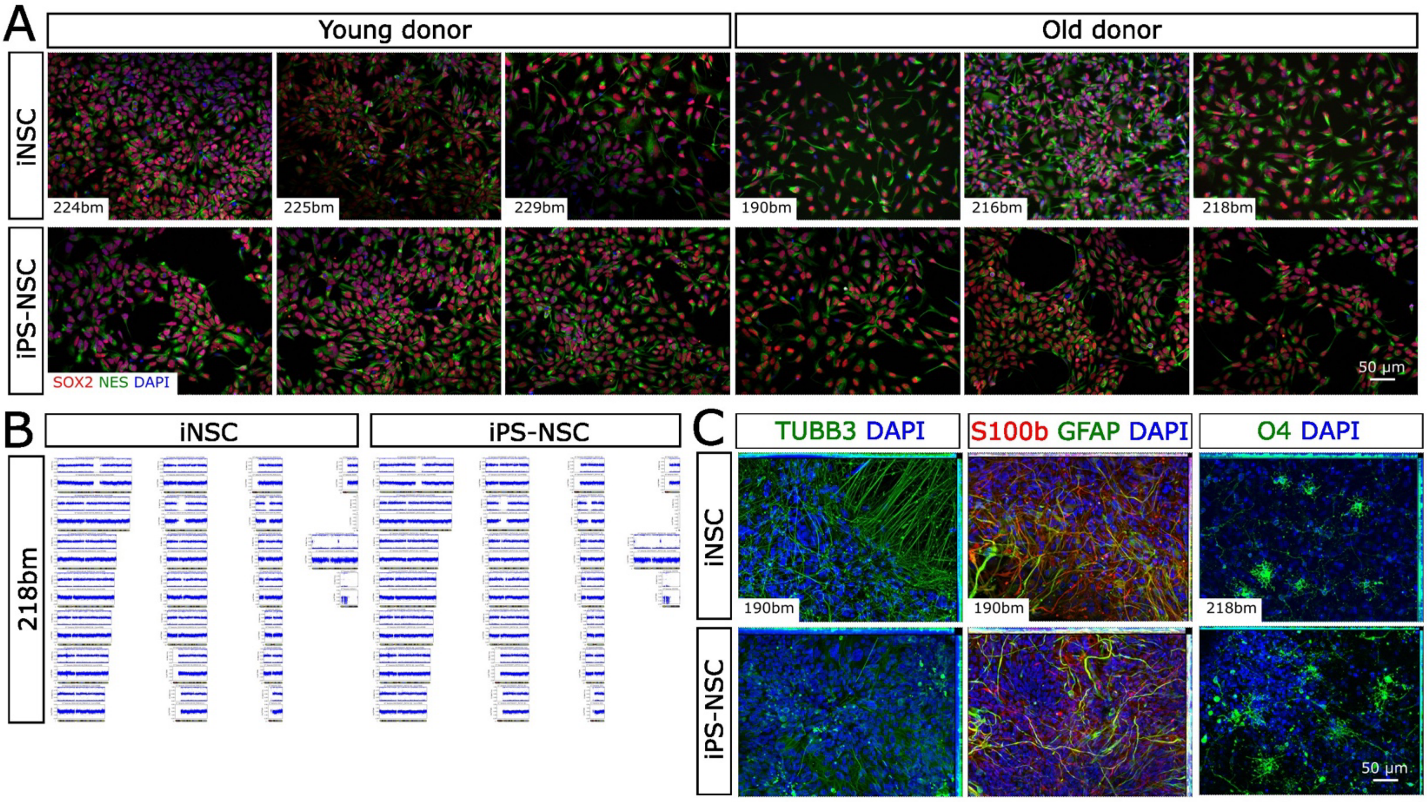


**Supplementary Figure S1**: **Young and old donor-derived EPCs can be programmed into genomically stable NSCs with tripotent differentiation capacity. (A)** Representative immunofluorescence images of young and old donor-derived iNSCs and iPS-NSCs stained against the NSC markers SOX2 and NES. Scale bar = 50 µm. **(B)** Exemplary SNP profiles of an isogenic set of high passage iNSCs and iPS-NSCs derived from one old donor (101 years of age). **(C)** Representative immunofluorescence images showing that old donor-derived iNSCs and isogenic iPS-NSCs can give rise to TUBB3-positive neurons, S100β- and GFAP-positive astrocytes as well as O4-positive oligodendrocytes upon differentiation. Scale bar = 50 µm.

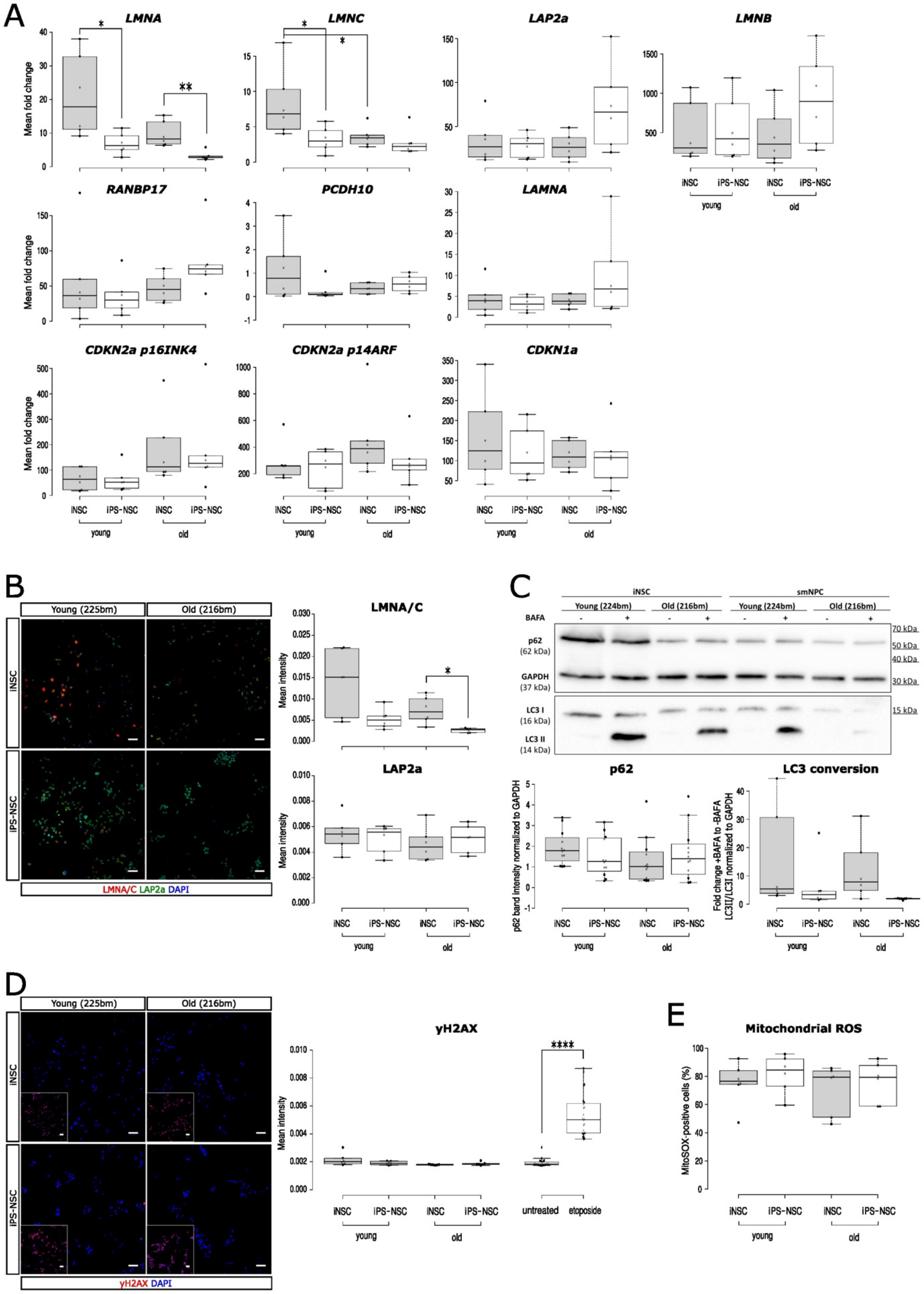


**Supplementary Figure S2: Cellular aging hallmarks are overall very similarly represented in young and old donor-derived iNSCs and iPS-NSCs.**

(Figure legend continued on the next page)

**(A)** Expression levels of the nuclear lamina-associated genes *LMNA*, *LMNC, LAP2$\alpha$* and *LMNB,* the three age-associated genes *RANBP17*, *LAMNA* and *PCDH10* (Mertens et al., 2015), and the two *CDKN2a* isoforms encoding for p16$^{INK4A}$ and p14$^{ARF}$ as well as *CDKN1a* encoding for p21 as measured by qPCR. N = 6 with three independent replicates of two genotypes per age group. Results of the Wilcoxon rank sum Kruskal Wallis post-hoc test: *LMNA* young donor-derived iNSCs vs. iPS-NSCs: $p$ = 0.03; *LMNA* old donor-derived iNSCs vs. iPS-NSCs: $p$ = 0.007; *LMNC* young donor-derived iNSCs vs. iPS-NSCs: $p$ = 0.03; *LMNC* young vs. old donor-derived iNSCs: $p$ = 0.03. **(B)** Left: Representative stainings for LMNA/C and LAP2a of iNSCs and iPS-NSCs of one genotype per age group. Scale bars = 50 µm. Right: CellProfiler-based quantification of nuclear LMNA/C and LAP2a immunofluorescence intensity. N = 6 with 10-24 pictures per three independent replicates of two genotypes per age group. Results of the Games-Howell ANOVA post-hoc test: LMNA/C old donor-derived iNSCs vs. iPS-NSCs: $p$ = 0.046. **(C)** Representative Western blot detecting p62, the LC3 isoforms I and II as well as the house-keeping protein GAPDH in iNSCs and iPS-NSCs under ± BAFA treatment conditions. Due to the number of samples, each biological replicate was run on two separate gels, with one genotype of each age group per gel. Underneath, the results of Western blot quantification are provided. N = 6 with three independent replicates of two genotypes per age group. For p62, both ± BAFA samples were considered. **(D)** Left: Representative stainings for γH2AX of iNSCs and iPS-NSCs of one genotype per age group. Inserts on the lower left show the etoposide-treated positive control of each condition. Scale bars = 50 µm. Right: CellProfiler-based quantification of nuclear γH2AX immunofluorescence intensity. N = 6 with 5-12 pictures per treatment condition in three independent replicates of two genotypes per age group. To compare untreated and etoposide-treated cells, data for all genotypes and replicates were pooled. Result of the Wilcoxon rank sum exact test: Untreated vs. etoposide-treated: $p$ = $6.202 \times 10^{-14}$. **(E)** Boxplot depicting the percentage of MitoSOX-positive cells according to flow cytometry analysis.

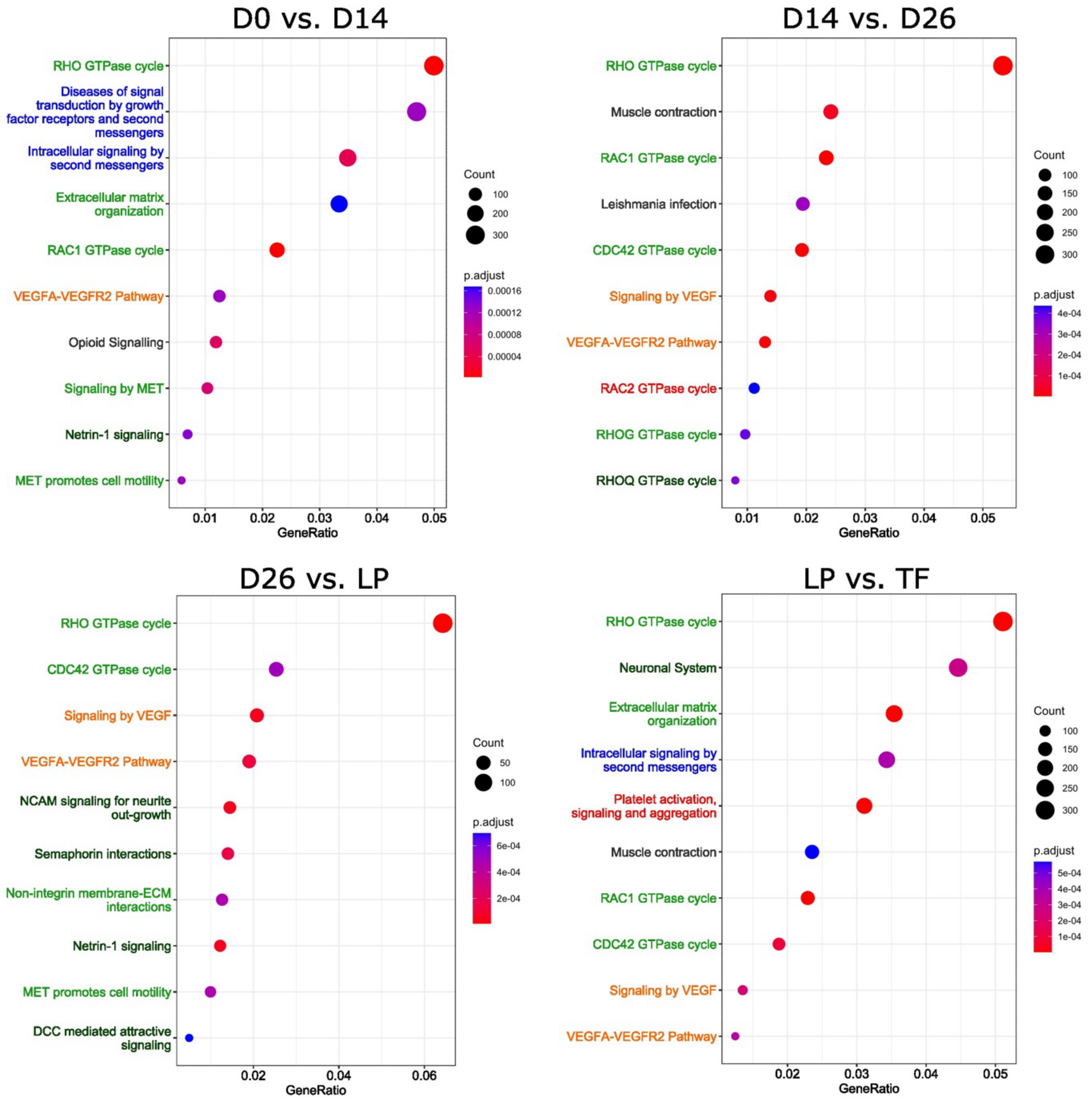


**Supplementary Figure S3**: **Global DNAm changes across all stages of iNSC conversion.** Dot plots representing the top 10 results of the Reactome pathway analysis performed on the mapped genes that were associated to the significant MVPs comparing day 0 to day 14 of conversion, day 14 to day 26 of conversion, day 26 of conversion to LP and LP to TF. Most pathways could be assigned to one of five categories: (i) neuronal system, especially neuronal migration, neurite outgrowth and pathfinding (dark green); (ii) cell migration, adhesion and division in multiple tissues, including pathways relevant to ECM and MET (light green); (iii) signal transduction (blue), (iv) angiogenesis (orange) and (v) the hematopoietic system (red).

### Supplementary Table

**Supplementary Table S1: Donors enrolled in this study.** * Blood sample from donor 82 was re-recruited one year after blood sample used for iNSC conversion was collected. ** iPS-NSC lines generated in the context of Sheng et al. M = Male; F = Female; N = No; Y = Yes.

| Donor code | Gender | Age in years | Included in Sheng et al. | iPS-NSCs generated | DNAm available | RNAseq available |
|---|---|---|---|---|---|---|
| 224 | M | 0 | N | Y | Y | Y |
| 225 | M | 0 | N | Y | Y | Y |
| 229 | M | 0 | N | Y | Y | Y |
| 108 | F | 31 | Y | N | N | Y |
| 96 | M | 33 | Y | Y** | Y | N |
| 107 | M | 35 | Y | Y** | Y | Y |
| 82 | F | 35/36* | Y | N | N | Y |
| 33 | F | 49 | Y | Y** | Y | Y |
| 190 | M | 50 | N | Y | Y | Y |
| 109 | M | 62 | Y | N | N | Y |
| 287 | F | 81 | N | N | Y | Y |
| 288 | M | 81 | N | N | Y | Y |
| 286 | F | 86 | N | N | Y | Y |
| 216 | M | 87 | N | Y | Y | Y |
| 218 | M | 101 | N | Y | Y | Y |

**Supplementary Table S2: GO terms that are commonly altered across all stages of iNSC conversion according to DNAm analysis.** Neural system-associated GO terms are highlighted in light grey. BP = biological process; CC = cellular component; DE = number of DEGs within this GO ID; MF = molecular function; N = number of all genes within this GO ID; P.DE = significance of GO term.

| ID | Ontology | Term | N | DE | P.DE |
|---|---|---|---|---|---|
| GO:0070161 | CC | anchoring junction | 821 | 632 | 1.85E-16 |
| GO:0042995 | CC | cell projection | 2252 | 1581 | 7.76E-15 |
| GO:0008092 | MF | cytoskeletal protein binding | 969 | 726 | 2.08E-14 |
| GO:0030036 | BP | actin cytoskeleton organization | 690 | 533 | 2.17E-14 |
| GO:0006928 | BP | movement of cell or subcellular component | 2182 | 1497 | 5.08E-14 |
| GO:0120025 | CC | plasma membrane bounded cell projection | 2152 | 1509 | 5.41E-14 |
| GO:0030054 | CC | cell junction | 2031 | 1443 | 1.28E-13 |
| GO:0007154 | BP | cell communication | 6470 | 4079 | 2.37E-13 |
| GO:0000902 | BP | cell morphogenesis | 1003 | 756 | 5.3E-13 |
| GO:0023052 | BP | signaling | 6457 | 4069 | 5.6E-13 |
| GO:0030029 | BP | actin filament-based process | 787 | 594 | 6.35E-13 |
| GO:0120036 | BP | plasma membrane bounded cell projection organization | 1525 | 1097 | 1.58E-12 |
| GO:0030030 | BP | cell projection organization | 1565 | 1123 | 1.7E-12 |
| GO:0032989 | BP | cellular component morphogenesis | 766 | 586 | 6.07E-12 |
| GO:0023051 | BP | regulation of signaling | 3452 | 2284 | 3.53E-11 |
| GO:0010646 | BP | regulation of cell communication | 3416 | 2260 | 3.61E-11 |
| GO:0040011 | BP | locomotion | 1907 | 1290 | 4.02E-11 |
| GO:0032502 | BP | developmental process | 6407 | 4074 | 4.23E-11 |
| GO:0048856 | BP | anatomical structure development | 5909 | 3782 | 4.39E-11 |
| GO:0007165 | BP | signal transduction | 5969 | 3729 | 5E-11 |
| GO:0048731 | BP | system development | 4855 | 3129 | 2.52E-10 |
| GO:0031175 | BP | neuron projection development | 976 | 719 | 4.69E-10 |
| GO:0007275 | BP | multicellular organism development | 5420 | 3468 | 5.39E-10 |
| GO:0009653 | BP | anatomical structure morphogenesis | 2685 | 1812 | 5.7E-10 |
| GO:0048666 | BP | neuron development | 1108 | 806 | 6.98E-10 |
| GO:0007010 | BP | cytoskeleton organization | 1376 | 965 | 7.4E-10 |
| GO:0022008 | BP | neurogenesis | 1611 | 1135 | 1.88E-09 |
| GO:0009966 | BP | regulation of signal transduction | 3053 | 2002 | 2.44E-09 |
| GO:0032990 | BP | cell part morphogenesis | 679 | 514 | 5.88E-09 |
| GO:0120039 | BP | plasma membrane bounded cell projection morphogenesis | 659 | 501 | 6.1E-09 |
| GO:0048858 | BP | cell projection morphogenesis | 663 | 503 | 8.26E-09 |
| GO:0048699 | BP | generation of neurons | 1499 | 1055 | 1.47E-08 |
| GO:0043005 | CC | neuron projection | 1313 | 923 | 1.89E-08 |
| GO:0050793 | BP | regulation of developmental process | 2494 | 1642 | 3.02E-08 |
| GO:0048812 | BP | neuron projection morphogenesis | 645 | 487 | 3.81E-08 |
| GO:0030182 | BP | neuron differentiation | 1356 | 954 | 1.32E-07 |
| GO:0007399 | BP | nervous system development | 2346 | 1592 | 1.67E-07 |
| GO:0061564 | BP | axon development | 506 | 379 | 3.17E-06 |
| GO:0048667 | BP | cell morphogenesis involved in neuron differentiation | 579 | 432 | 4.77E-06 |
| GO:0007409 | BP | axonogenesis | 460 | 348 | 5.98E-06 |
| GO:0000904 | BP | cell morphogenesis involved in differentiation | 723 | 528 | 1.07E-05 |
| GO:0034330 | BP | cell junction organization | 699 | 501 | 2.04E-05 |
| GO:0030424 | CC | axon | 619 | 444 | 8.78E-05 |
| GO:0007411 | BP | axon guidance | 277 | 212 | 0.000364904 |
| GO:0097485 | BP | neuron projection guidance | 278 | 212 | 0.000525281 |
